## Supplementary Material for "Glycolipid recognition and binding by Siglec-6 hinges on interactions with the cell membrane"

**Assessing the potential involvement of non-canonical Arg residues in the V-set domain of Siglec-6 in the recognition and binding of GM1os.** Previous work by some of us^1^ clearly showed that Siglec-6 binds gangliosides with a mechanism independent of the canonical Arg, namely Arg122. Due to the presence of multiple Arg and Lys residues in the V-set domain of Siglec-6, it is reasonable to hypothesize that one of those may have replaced or could complement the function of the conserved Arg, as some of us recently demonstrated in the case of Siglec-10^2^. To address this matter we ran a set of three uncorrelated molecular dynamics (MD) simulations of the non-glycosylated Siglec-6 (see results in the main text) in an equilibrated water box, in the presence of seven GM1os molecules. The starting structure in all replicas was built with one of the GM1os bound to the canonical Arg122 to occupy the canonical site, while another was positioned in proximity to Arg92, , see **Figure S.1,** which has been previously highlighted as a potential (non-canonical) binding Arg^1^. We focused our attention to the Arg in the V-set domain, but monitored contacts with all Arg on the Siglec-6 accessible surface, as these could be involved in both *cis,* and *trans* binding events. The results show that none of the GM1os, except the one bonded to the canonical Arg122, engage in stable interactions, see **Figure S.1**. The Arg92 is engaged in a stable salt bridge with Asp115, which is never disrupted during any of the three independent MD simulations. As a caveat, it is important to underline that additive force fields, such as the one we used to run the MD simulations in this work, are notoriously biased towards enhancing electrostatic contacts^3^ and thus tend to overestimate the stability of already strong hydrogen bonding interactions, such as salt bridges^4^. In this specific case an analysis of the Siglec-6 3D structure indicates that such salt bridge interaction may be essential to the correct folding of the V-set domain, and mutation of Arg92 (or of Asp115) would compromise the folding, which explains the low expression levels of the R92A mutant reported in earlier work^1^.

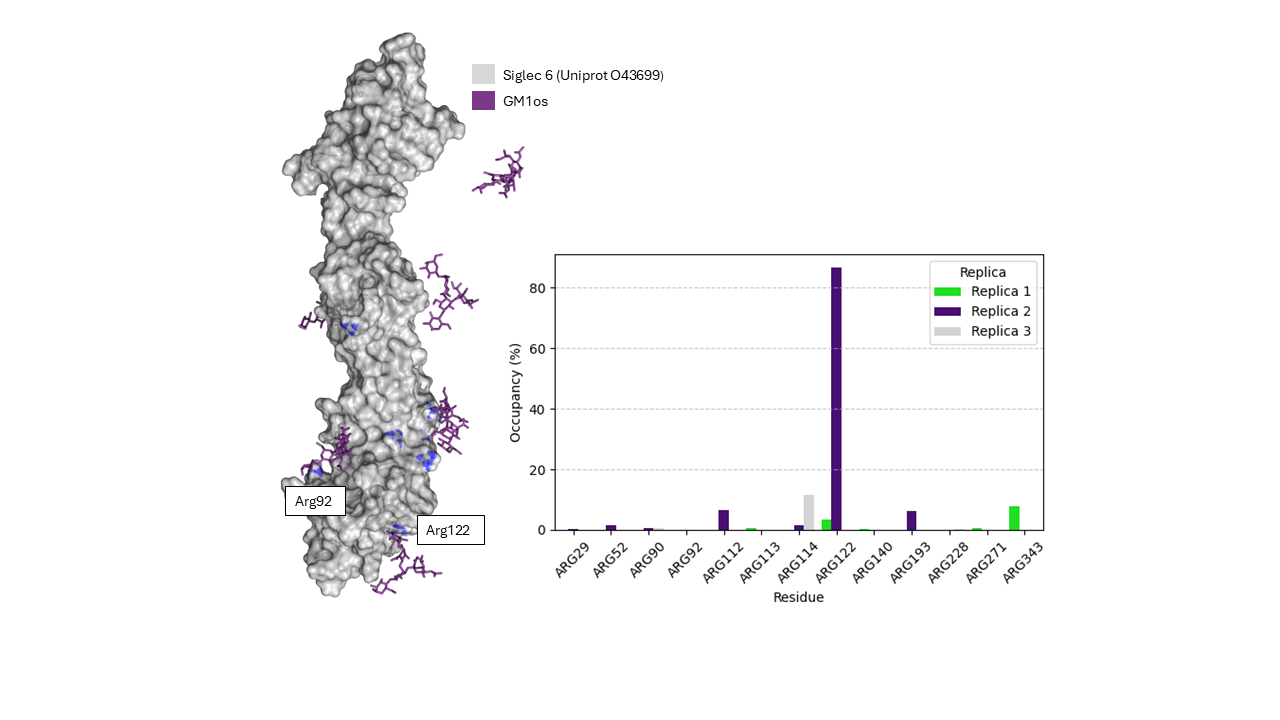

**Figure S.1. Left:** 3D surface representation of Siglec-6 (AF-O43699, gray) with seven GM1os molecules (purple sticks), including one placed at the canonical sialic acid binding site (Arg122) and others distributed in possible non canonical regions of the protein. **Right:** Bar plot showing the occupancy of GM1os interactions with selected ARG residues across three independent MD replicas. Occupancy is assessed in terms of distance between guanidinium C (CZ) of Arg and the carboxyl carbon (C1) of the Neu5Ac with a threshold of 5 Å for binding. Molecular rendering with VMD (<https://www.ks.uiuc.edu/Research/vmd/>) and bar plot with MS Excel.

**MD simulations of the fully *N*-glycosylated Siglec-6 in complex with membrane bound GM1.** We ran two independent MD simulations of 500 ns each on a 3D model of the fully glycosylated Siglec-6 in complex with an isolated GM1 ganglioside embedded in the phospholipid bilayer. The molecular composition of the 3D model is described in the Methods section and, aside from the N-glycans, is analogous as the one presented in the main text. The Siglec-6 sequence (453 aa) carries seven *N-*glycosylation sequons, with only one in the V-set domain at Asn103, see **Figure S.2**. All sites are occupied with a biantennary monogalactosylated complex *N*-glycan (GlyTouCan ID G99129GB; 1478.5 Da). As represented in the snapshot from the MD simulations shown in **Figure S.2**, only the *N*-glycans at N104 and N149 interact with the protein, and more specifically engage in contacts that may affect the relative orientation and dynamics of the V-set domain (d1) and the adjacent C2 domain (d2). The RMSD values calculated over the entire extracellular region of the glycosylated system (i.e. over the backbone atoms of d1, d2, and d3) are higher than the non-glycosylated one, see **Figure S.2.d**. This increase is not due to destabilization by glycans, but rather reflects the higher conformational mobility of the distal d3 domain, where glycans such as those at N258 and N295 are not involved in protein contacts and remain solvent exposed and dynamic. It is important to note that d3 is connected to the transmembrane domain, and in a complete cellular context, its dynamics may be constrained by interactions with the membrane. Instead, our data support a stabilizing function of the glycans at N103 and N149, which maintain the relative positioning and rigidity of domains d1 and d2. When compared to the non-glycosylated Siglec-6/GM1 complex, the fully glycosylated Siglec-6 shows a slightly lower average RMSD value relative to the AF (AF-O43699-F1) backbone, suggesting a reduced interdomain flexibility and higher structural stability of the d1–d2 interface, see **Figure S.2.c**. However, the relative flexibility of the system demonstrated by the spread around the median RMSD value in the KDE plots in **Figure S.2.d** for both glycosylated and non-glycosylated systems, precludes defining any clear role of the *N*-glycans in affecting the rigidity of the V-set domain relative to the adjacent C2-domain.

As an important note, the binding to the GM1 is not affected by the *N*-glycosylation neither directly, nor indirectly. This is further supported by the MD simulations of the fully glycosylated Siglec-6/GM1 complex, which consistently show stable anchoring on the membrane through the interactions with Trp127 and Lys126. In the first MD simulation (MD1), the salt bridge between Arg with Neu5Ac was restrained for the first 300 ns. When restraints were released, the system remained stable and preserved the salt bridge and the membrane contacts for the remaining simulation time. In the second MD simulation (MD2), where no restraints were applied, the Arg and Neu5Ac interaction was lost immediately, yet the system remained stable over 500 ns. This stability was maintained through persistent interactions between terminal Gal of GM1 and Asp70 residue in the C-C' loop, as well as the continued involvement of Lys126 and Trp127 in membrane interaction.

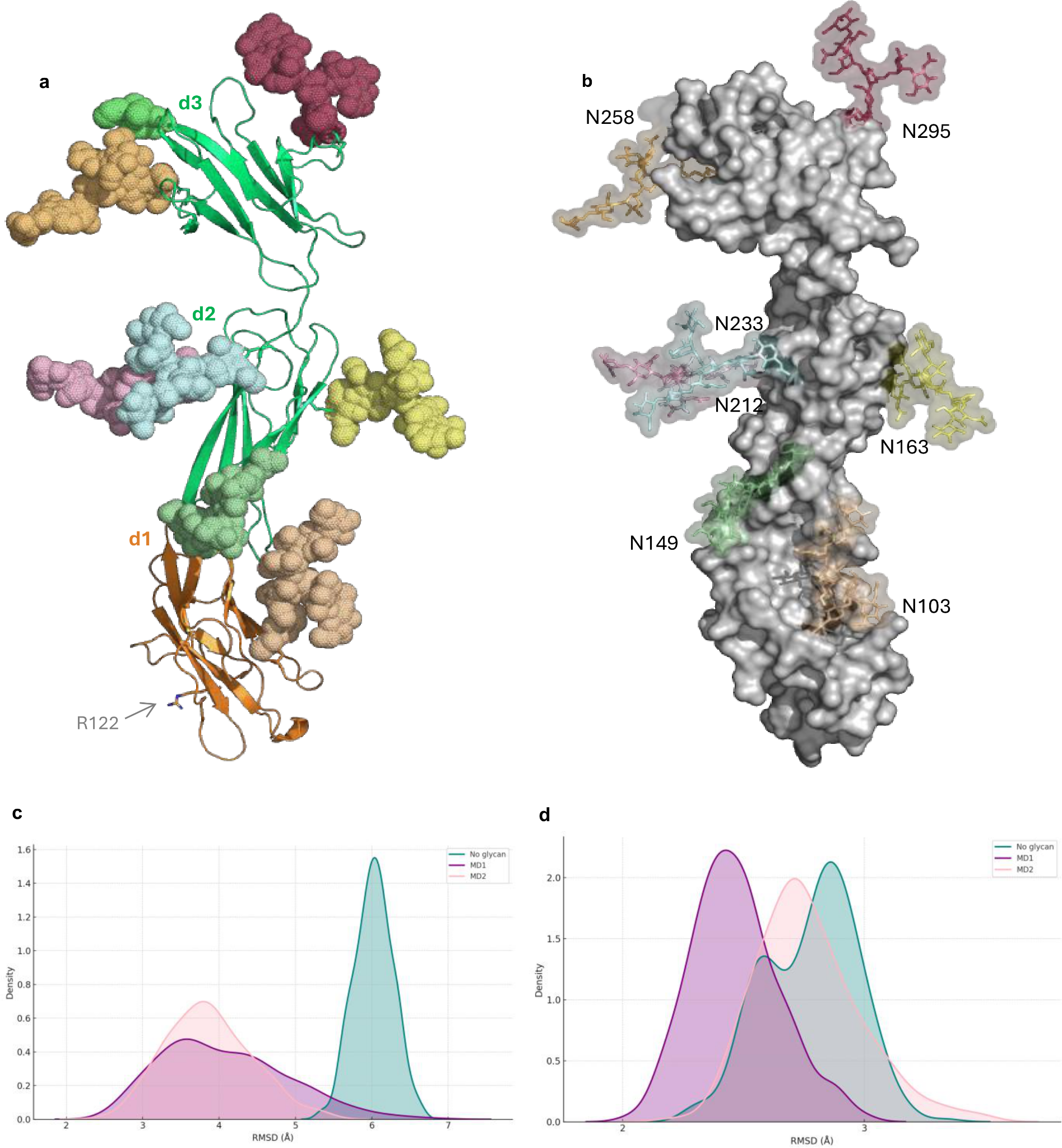

**Figure S.2. a.** 3D structure of the Siglec-6 extracellular region from AF model AF-O43699-F1, shown as cartoons. The terminal V-set binding domain (d1) is shown in orange and the C2-set domains (d2 and d3) are shown in green. *N*-linked glycans are rendered with atoms as vdW spheres and color-coded individually. The seven *N*-glycosylation sequons were assessed as occupied with the GlcNAc Scanning tool in GlycoShape ReGlyco^5^. **b.** Representative snapshot of the Siglec-6 from the MD simulation with the N-glycosylation sites labelled. **c.** Kernel density estimates (KDE) distributions of the backbone RMSD values along the trajectory, corresponding to the Siglec-6 calculated for d1 and d2. The results corresponding to the non-glycosylated Siglec-6 are shown in green and for the fully glycosylated Siglec-6 in purple and pink for MD1 and MD2, respectively). **d.** Kernel density estimates (KDE) distributions of the backbone RMSD values calculated along the MD trajectories for the backbone of the Siglec-6 extracellular region, i.e. d1 to d3. Molecular rendering with *pymol* ([www.pymol.org](http://www.pymol.org)) and KDE plots with *seaborn* (<https://seaborn.pydata.org/>).

**MATERIALS AND METHODS**

**Computational Methods**

**MD simulations of Siglec-6 with multiple GM1os.** The 3D structure of Siglec-6 was obtained from the AlphaFold database (AF-O43699-F1) ^6,7^, and the structures of GM1os were retrieved from the GlycoShape database ^5^. One GM1os molecule was manually placed at the canonical sialic acid-binding site near Arg122, while six additional GM1os molecules were randomly distributed around the protein surface to investigate for potential alternative binding sites. The protein was parameterized using the AMBER ff14SB force field, and the glycans with GLYCAM06j-1. Simulations were performed using AMBER18 . Following 500,000 steps of steepest descent minimization, the system was heated in two stages (0–100 K and 100–300 K, 500 ps each) in the NVT ensemble using Langevin dynamics (γ_ln = 1.0 ps⁻¹). Equilibration was carried out in the NPT ensemble for 500 ps at 1 atm using a Berendsen barostat. All restraints on the protein heavy atoms were then removed, and three independent 300 ns production runs were performed from different initial velocities. Due to intrinsic limitations of the force field in modeling systems with multiple free glycans, the simulations were halted prematurely ^3^. Residue specific occupancies were calculated by monitoring persistent contacts (within 5 Å) between the carboxyl group of the sialic acid and the guanidinium group of ARG side chains over time.

**Modelling of the glycosylated Siglec-6**

The AlphaFold-predicted structure of Siglec-6 (AF-O43699-F1) was scanned using GlcNAc Scan tool^5^, which identified seven putative *N*-glycosylation sites across the extracellular region. The glycan 3D structures were retrieved from the GlycoShape database and added to each predicted sequon using Re-Glyco, which optimizes the orientation of the attached N-glycans (GlyTouCan ID: G99129GB) to minimize steric clashes and ensure proper linkage geometry. The glycosylated PDB output from Re-Glyco was then used as input in CHARMM-GUI Membrane Builder to generate the membrane system. MD simulations were performed as described in the Methods section “MD simulations of GM1 and GM2 in complex with Siglec-6”.

**Table S.1: Primers used in this work.**

| **Primer Name** | **Sequence** |
| --- | --- |
| Siglec-6 Fwd | AGC AGC GCT AGC ATG CAG GGA GCC CAG GAA GCC |
| Siglec-6 Rvs | AGC AGC ACC GGT TCA CTT GTG TAT CTT GAT TTC |
| Sig-6 K124A Fwd | CGG TTG AAG TCC GCA TGG ATG AAA TAC |
| Sig-6 K124A Rvs | GTA TTT CAT CCA TGC GGA CTT CAA CCG |
| Sig-6 Rvs 3D | AGC AGC ACC GGT CCT GCC TTC TGG TTT CCA ATG |
| Sig-6 K126A Fwd | CGG TTG AAG TCC GCG TGG ATG AAA TAC GG |
| Sig-6 K126A Rvs | CCG TAT TTC ATC CAC GCG GAC TTC AAC CG |
| Sig-6 W127A Fwd | GTT GAA GTC CAA AGC CAT GAA ATA CGG TTA TAC |
| Sig-6 W127A Rvs | GTA TAA CCG TAT TTC ATG GCT TTG GAC TTC AAC |
| Sig-6 K129A Fwd | GTC CAA ATG GAT GGC ATA CGG TTA TAC |
| Sig-6 K129A Rvs | GTA TAA CCG TAT GCC ATC CAT TTG GAC |

**Cell culture.** Flp-In Chinese hamster ovary (CHO) cells were cultured as previously described^1^, but in short, CHO cells were cultured in DMEM/F12 Media (Gibco) supplemented with 5% (V/V) fetal bovine serum (Gibco), penicillin (100 U/mL), and streptomycin (100 μg/mL). Cells were grown at 37 °C and 5% CO_2_ in tissue culture dishes.

**Stable transfection of Flp-In CHO Cells*.*** Flp-In Chinese^1^.

**Liposomes preparation**. Liposomes were prepared by hydrating lipid thin consisting of DSPC (61.5 mol%), cholesterol (38 mol%), and PEG_45_-DSPE (0.4 mol) and AF647-PEG_45_-DSPE films with PBS pH 7.4. Glycolipids were added at the expense of DSPC. The lipid solutions were extruded using an Avanti-mini extruder using 800 nm and 100 nm filters respectively. Liposomes were then validated with dynamic light scattering. More details regarding liposome production can be found elsewhere^1^,^15^.

**Siglec-Fc expression and purification.** The procedure for Siglec-Fc expression and purification can be found elsewhere^16^)

**Cell assay**. 150,000 cells were added to a 96 well U-bottom microplate. Liposome solutions were prepared at 50 μM in 1% (g:mL) BSA PBS pH 7.4. 50 μL of liposome solution was added to the cells and the cells were incubated with the liposomes for 30 min at 37 °C. The cells were then washed with PBS twice. Following the washes, the cells were then resuspended in anti-Siglec antibody solution (1/250 V:V antibody in 500 µM EDTA, 1% BSA (g:mL)). The cells were then incubated at 4 °C for 30 min. The cells were again washed and then analyzed by flow cytometry. Antibody information can be found in Table S.2, and general gating strategy can be found in Figure S.3.

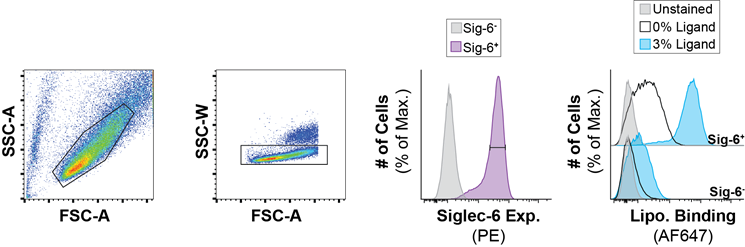

**Figure S.3.** Gating strategy used to measure glycolipid liposome binding to Chinese hamster ovary cells expressing Siglec-6 mutants. Liposome binding was measured using 3 mol% compound NGL1 ^1^.

Chemical structure of neoglycolipid ligand (NGL1) used in this experiment. Synthesized as described previously ^1^.

**Table S.2:** Antibodies used in this work

| **Antibody** | **Supplier** | **Cat. No.** | **Label** | **Clone** | **Isotype** | **Dilution** |
| --- | --- | --- | --- | --- | --- | --- |
| anti-Siglec-6 | R&D Systems | FAB2859P | PE | 767329 | Mouse IgG2A | 1:250 (V:V) |

**Flow cytometry.** Flow cytometry measurements were collected on a 5-laser Fortessa X-20 (BD Bioscience). All the resulting data were analyzed using FlowJo (10.5.3) software (BD Biosciences)

**Siglec-Fc ELISA.** A detailed description of the approach can be found elsewhere ^1,15^. In short, 50 μL of 50 μM glycolpid solution was transferred to a 96 well ELISA microplate. The plate was left at 37 °C for 2 to remove the ethanol. The wells were blocked with 5% (g:mL) bovin serum albumin (BSA) for 1 h at room temperature. During the blocking step, the Siglec-Fc-Strep-Tactin solution (Siglec-Fc solution: 2 μg/mL, Strep-Tactin-HRP: 0.13 μg/mL-IBA) was prepared and complexed at room temperature for at least 30 min at room temperature. The blocking buffer was then removed, and the Siglec-Fc solution was added for 2 h at room temperature. Unbound Siglec-Fc solution was then removed, and the amount of binding was measured using TMB substrate. The reaction was then quenched using 1 M H_3_PO_4_ and the Abs_450_ was measured using a Molecular Devices SpectraMAX iD5.
